# Epigenetic Clocks Reveal Age Acceleration and Shared Methylation Remodeling Across Cancers

**DOI:** 10.64898/2026.08.27.747695

**Authors:** Saleh Sereshki, Stefano Lonardi

## Abstract

DNA methylation-based epigenetic clocks estimate biological age from methylation profiles, and the difference between predicted biological age and chronological age is commonly described as age acceleration (AA). We compared AA across eight cancer types—lung, colorectal, breast, thyroid, bone marrow and blood, kidney, uterus, and head and neck—using seven epigenetic clocks and 5,528 publicly available samples. Across the 56 cancer type–clock combinations, tumor tissues showed higher average AA than normal tissues in 44 comparisons. The uterus cohort showed the clearest deviation from this overall trend, with normal samples exhibiting higher AA for six of seven clocks. Analyses of paired normal and tumor samples generally showed higher predicted ages and greater variability in tumor samples. We additionally examined age-associated methylation changes and the ability of clock CpGs to distinguish tumor from normal tissue. Several discriminatory CpGs were shared across cancer types and frequently showed tumor-associated hypermethylation at cancer-related loci. Small subsets of top-ranked CpGs captured substantial discriminatory information. Age-stratified subsampling preserved the main AA patterns, suggesting that chronological-age differences did not explain the observed tumor–normal differences. Overall, these findings highlight broad cancer-associated alterations in epigenetic aging together with substantial cancer type- and clock-specific heterogeneity.

## 1 Introduction

Biological age is a latent value that reflects the extent of aging-driven biological changes in an organism. Since DNA methylation undergoes age-related changes across various tissue types, it can be used as a biomarker for predicting biological age, the so-called *DNAm* age. [1, 2, 3, 4, 5]. Various DNAm age predictors have been developed (called *epigenetic clocks*) to estimate the biological age based on DNA methylation profiles (see, e.g., [2, 6, 7, 8, 9, 10]). These clocks differ based on the tissue specificity, the process of CpG site selection, and the methodological approach used. For instance, the Horvath clock employs a penalized regression model to estimate chronological age by leveraging DNA methylation data from 353 CpG sites, making it applicable across a variety of human tissues [2]. In contrast, the Hannum clock is designed for blood samples and predicts biological age using 71 CpG sites [6]. The Levine clock uses CpG sites that are not only predictive of chronological age but also highly informative regarding lifespan and health span, thereby integrating measures associated with age-related health outcomes and mortality [7]. Very often there are difference between an the biological age (DNAm) and chronological age of an individual. The difference between DNAm age and chronological age is known as *age acceleration* (AA). A slower biological aging has been shown to be associated with longevity and lower risk of age-related disease [11]. A faster biological aging has been linked to cardiovascular disease, diabetes, psychiatric disorders and cancer [12, 13, 14, 15, 16]. Both cancer and aging result from cumulative cellular damage and share similar underlying mechanisms [17, 18]. These common dysregulations extend to epigenetic changes and DNA methylation, contributing to AA in cancer patients [19, 20]. Numerous studies have demonstrated that AA can serve as a biomarker for early cancer detection and has the potential to be used in cancer therapeutics [21, 22, 23, 24, 25, 26].

DNA methylation patterns and its cancer-associated alterations are tissue-specific [27, 28, 29]. This specificity should be considered when using AA in clinical applications. For example, a recent study by *Hao et al*. [30] indicates that in uterine cancer, AA is associated with cancer development, whereas the opposite is observed in other cancer types, such as head/neck, colorectal, and breast cancers [31, 32, 33, 34]. In another study, Dugué *et al*. employed three epigenetic clocks to assess AA as a prognostic biomarker for seven cancer types [35]. The authors found a significant association between AA and cancer risk for certain cancer types, including lung, kidney, and colorectal cancers, but observed limited evidence for prostate cancer. Studies that did not find strong evidence on the role of AA in cancer typically used only a single clock or a limited set of clocks to measure AA, or they focused exclusively on AA in a single cancer type.

in this study, we investigate AA in eight different cancer types, namely (i) lung, (ii) colorectal, (iii) breast, (iv) thyroid, (v) bone marrow and blood, (vi) kidney, (vii) uterus, and (viii) head and neck, using publicly available datasets. We compared seven different epigenetic clocks to measure AA in a total of 5,528 samples (see Section 2.2 for details on the seven clocks). We analyzed age acceleration in individuals with data for both normal and tumor samples, as well as the distribution of AA within samples from each cancer type. We further investigated the methylation patterns of the CpGs used by these clocks, assessed the ability of individual CpGs to distinguish tumor from normal tissue, examined the extent to which discriminatory CpGs were shared across cancer types and their genomic annotations, and compared the tumor–normal discriminatory information captured by CpG sets from different clocks. To the best of our knowledge, this represents one of the most extensive comparative analyses of epigenetic age acceleration and clock-associated methylation patterns across multiple cancer types and epigenetic clocks. We envision that our findings will contribute to a better understanding of the relationship between epigenetic aging and cancer.

## 2 Materials and Methods

### 2.1 Data sets

All datasets analyzed in this study were obtained from The Cancer Genome Atlas (TCGA) via the Genomic Data Commons (GDC) Data Portal. For each cancer type, all available Illumina 450k methylation array datasets were downloaded. Metadata, including patient age and tissue types, were also retrieved from the GDC. Methylation levels for each cohort were then calculated using a custom Python script.

### 2.2 The Epigenetics Clocks

Seven epigenetic clocks were used in this study, namely 1) Horvath’s clock, one of the earliest and most widely used, which employs the methylation levels of 353 CpGs to predict age across various tissues; 2) Hannum’s clock, focused on blood samples, which utilizes the methylation levels of 71 CpGs; 3) Levine’s clock, also known as PhenoAge, which uses the methylation levels of 513 CpGs to predict age from blood samples; 4) BNN leverages Horvath’s clock with a Bayesian Neural Network for age prediction; 5) Horvath’s Skin+Blood clock targets specific tissues and uses the methylation levels of 391 CpGs; 6) the BLUP clock utilizes the methylation levels of 319,607 CpGs to predict age from blood and saliva samples and 7) the EN clock, which employs the methylation levels of 514 CpGs to estimate age from blood and saliva samples. The Horvath, Levine, Horvath Skin, and EN clocks utilize elastic net regression, while the Hannun clock relies on linear regression. The BLUP clock employs best linear unbiased prediction models.

The predicted ages of samples were determined using the ‘methyl-clock’ library in R, based on the DNA methylation levels obtained avove.

### 2.3 Data sub-sampling

To create a balanced subset for analysis, we implemented a data subsampling strategy that maintains a consistent ‘Tumor’ to ‘Normal’ ratio across all age groups while maximizing sample inclusion. We divided the dataset into 10-year age bins based on the chronological age. In each bin, we calculated the ratio of Tumor to Normal samples and identified the smallest ratio among all bins. Using this smallest ratio, we included all available Normal samples. We determined the number of Tumor samples to include by multiplying the Normal sample count by this ratio and rounding down to the nearest whole number. These Tumor samples were then randomly selected from all available Tumor samples.

### 2.4 CpG-Level Tumor–Normal Discrimination and Cross-Tissue Overlap

To characterize tumor-associated methylation patterns within the CpGs used by each epigenetic clock, we first visualized their methylation beta values in normal and tumor samples for each cancer type. Samples were separated by tissue status and ordered by chronological age within each group, while CpGs were ordered by their mean methylation level across normal samples. For each clock and cancer type, we then assessed the discriminatory contribution of individual CpGs using univariate logistic regression, with tumor status as the binary outcome and methylation beta value as the predictor. Statistical significance of the regression coefficient was evaluated using a two-sided Wald test, and multiple-testing correction was performed using the Benjamini–Hochberg procedure. CpGs with FDR < 0.05 were classified as hypermethylated or hypomethylated in tumors according to the sign of the regression coefficient. Exact intersections of significant CpGs across the eight cancer types were summarized using UpSet plots while preserving the direction of methylation change. For BNN, EN, Hannum, Horvath, Levine, and Skin Horvath, CpGs that remained significant with the same direction across all eight tissues were further annotated using the Illumina HumanMethylation450 manifest to identify associated genes, gene-region context, and CpG-island context. Their genomic distributions were compared descriptively with the distribution of all CpGs represented on the 450K array. The corresponding cross-tissue UpSet and genomic-context analyses for the individual clocks are shown in Supplementary Figures 6 and 7.

### 2.5 Elastic-Net Comparison of Clock-Specific Discriminatory Power

To compare the amount of tumor–normal discriminatory information captured by CpGs from different epigenetic clocks, CpGs were ranked separately for each clock and cancer type according to their univariate tumor–normal association P-values. Increasing subsets of the top 10, 20, 40, 80, 160, 320, and 640 CpGs were evaluated, where sufficient CpGs were available. For each subset, an elastic-net logistic regression classifier was fitted using methylation beta values as predictors, with missing values imputed using the median and features standardized before model fitting. Equal contributions of L1 and L2 regularization were used (l1-ratio = 0.5). Predictive performance was evaluated using five-fold stratified cross-validation, and out-of-fold probabilities from all folds were pooled to calculate the area under the receiver operating characteristic curve (ROC AUC) and the area under the precision–recall curve (PR AUC). ROC AUC was compared with the random-classification value of 0.5, whereas the baseline for PR AUC was defined by the proportion of tumor samples in each cohort.

## 3 Results

We analyzed eight cohorts, each representing a specific type of cancer (thyroid, colorectal, bone marrow and blood, uterus, head and neck, breast, kidney, and lung) with corresponding normal and tumor tissue samples (see Section 2.1 for details). Figure 1-A shows the age distribution for all normal and tumor samples. Figure 1-B illustrates the number of tumor and normal samples in each cohort.

**FIGURE 1.**
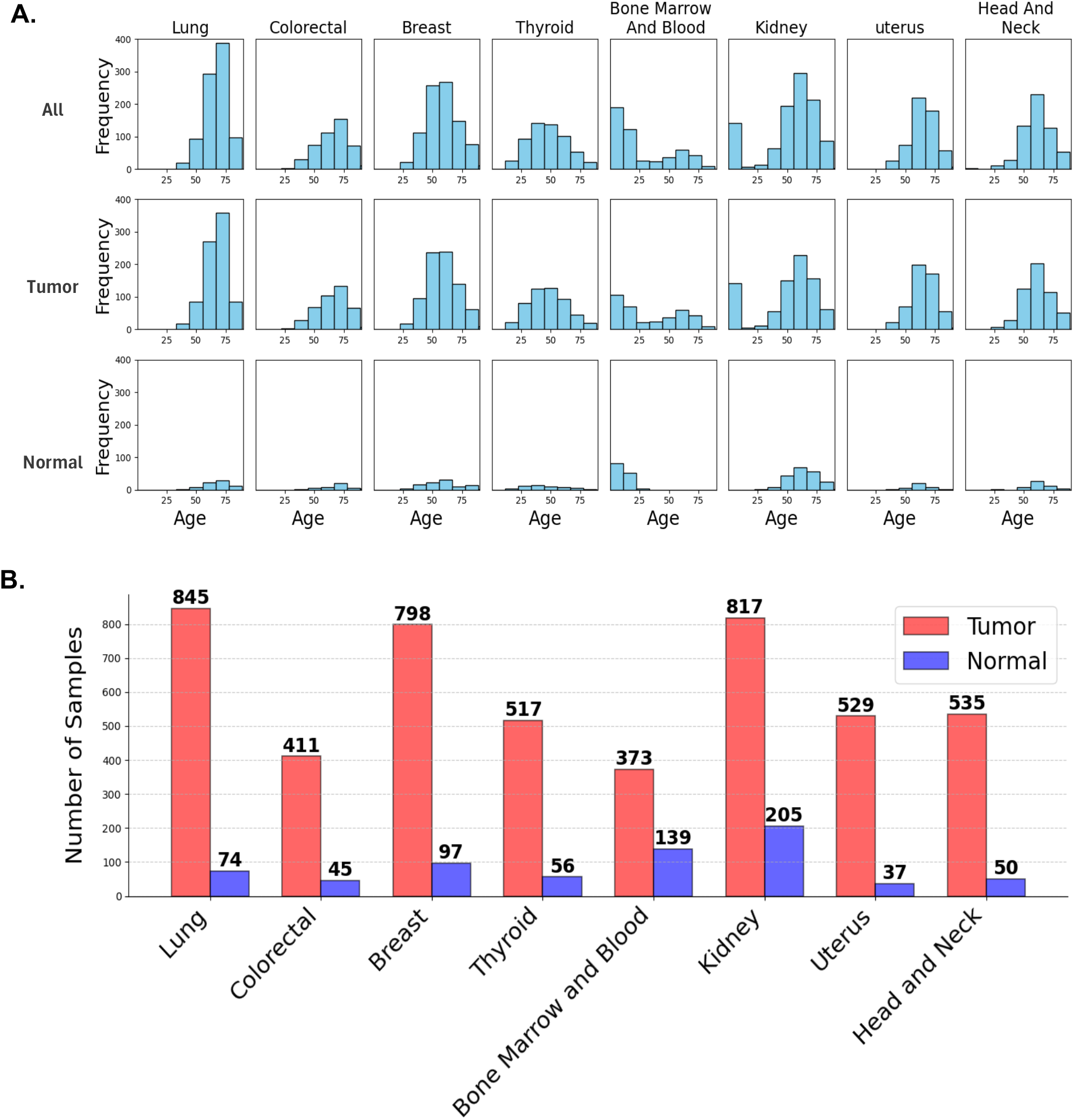
**A:** Age distribution of all tumor and normal samples, **B:** Distribution of normal and tumor samples in 8 cohorts

As said, we used seven epigenetic clocks to calculate the epigenetic age from the methylation data (see Section 2.2 for details). We determined AA by computing the difference between predicted and chronological age. Experimental results in Figures 2-A and B show two scatter plots, in which the x-axis is the average AA for a normal sample and the x-axis is the average AA for the corresponding tumor sample. Each point represents a specific cohort and clock method. Each plot shows a total 56 points, 8 cohorts × 7 clocks. In Figure 2-A, points are color-coded by their cohorts, whereas in Figure 2-B, points are color-coded by the methylation clock used.

**FIGURE 2.**
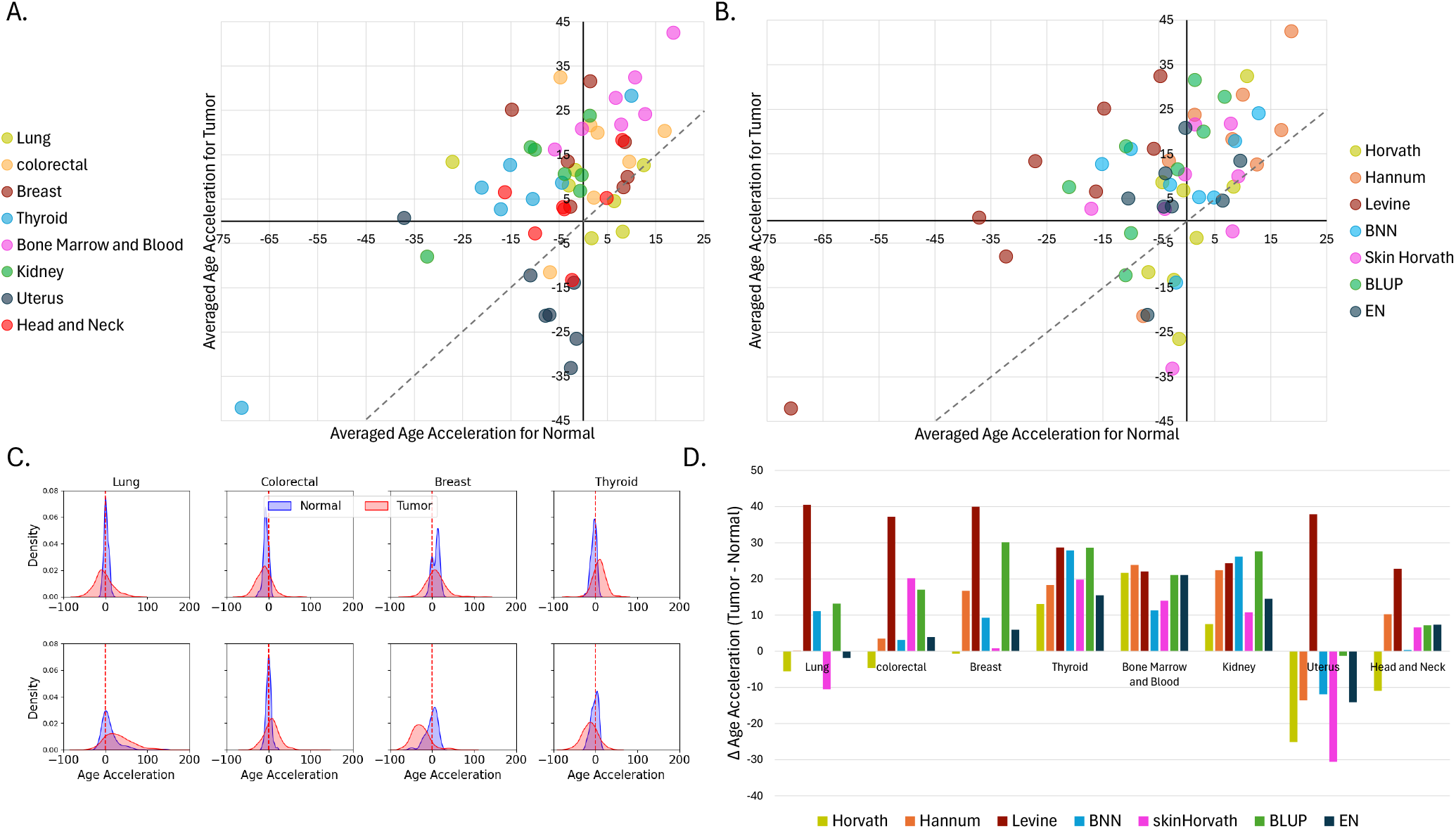
**A-B:** Comparing the averaged age acceleration (AA) in tumor versus normal samples across eight cancer types and seven epigenetic clocks; **A:** points are color-coded by cancer type; **B:** points are color-coded by the clock used; **C:** AA distribution of normal and tumor samples in eight cancer types (using Horvath clock); **D:** Difference in averaged AA between normal and tumor samples across cancer types and epigenetic clocks

**FIGURE 3.**
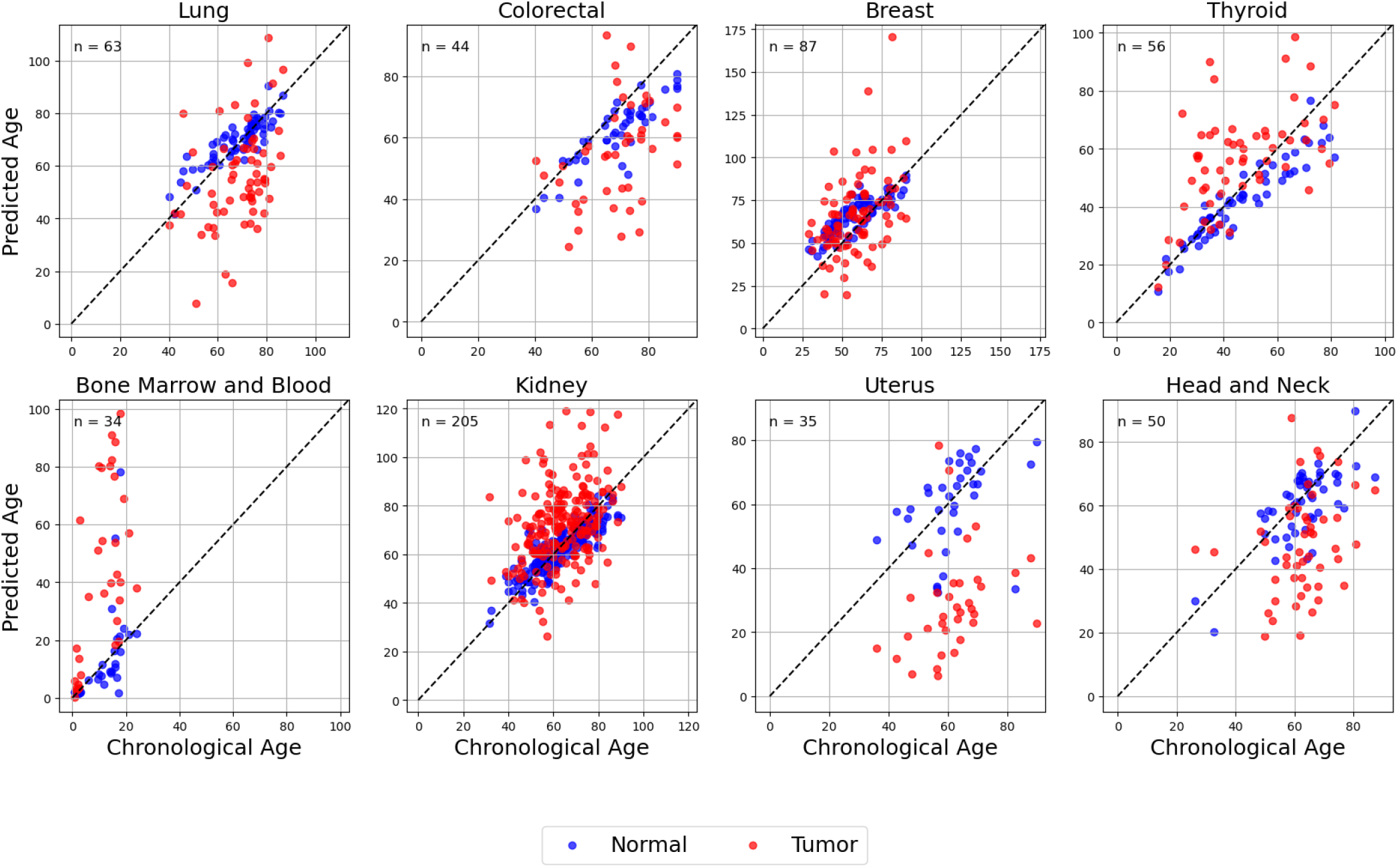
Comparing Horvath DNAm age and chronological age for normal and tumor samples across eight cancer types

Observe that in 44 cases out of 56, the points are above the diagonal, indicating that tumor tissues have higher average AA compared to normal tissues. Also observe that the uterus cohort deviates from this trend, with normal samples showing higher AA than tumor samples in six out of seven clocks. Additionally, the Horvath clock demonstrated higher AA in normal samples in 5 out of 8 cohorts, the highest frequency observed among all clocks evaluated. Overall these results indicates a strong association between cancer and AA, which is consistent with previous studies [31, 32, 33, 34].

In Figure 2-C we plotted the AA distribution of eight cancer types using Horvath clock. Observe that not only the average AA is different, but also the distribution. We also calculated the ΔAA by subtracting the average AA of normal samples from average AA of tumor samples, for each of the 56 clock-cohort pairs. Figure 2-D illustrates these differences across all cohorts and clocks. Observe that the ΔAA is positive for all types of cancer, except for the uterus cohort.

Supplementary Figure 1 shows the distribution of AA for each combination of clock and cohort. In all cases, a significant difference in distribution was observed between the AA of normal and tumor samples. Tumor samples exhibited a flatter distribution with a broader range of age acceleration, while normal samples were confined to a narrower range.

### 3.1 Analysis of Paired Samples

Next, we expanded our analysis to the individual level. In each cohort, we identified pairs of normal and tumor samples from the same individual. For each patient, we plotted the biological (predicted) age obtained from the seven epigenetic clocks and their actual chronological ages (see Supplementary Figure 2). Red points represent tumor samples, and blue points represent normal samples. Each individual is represented by two points in these scatter plots. Generally, tumor (red) samples exhibit greater dispersion in the plots, indicating that the DNAm clocks perform more variably for these samples. Moreover, in most cases (except for the uterus cohort) the predicted ages of tumor samples are higher than those of normal samples.

### 3.2 Controlling for Age Distribution

As shown above, the age distributions of normal and tumor samples was different. This could potentially bias our analysis because the chronological age could influence the magnitude of AA. To compensate for this effect, we implemented a subsampling strategy to ensure that normal and tumor samples had similar age distributions within 10-year age bins (see Methods for details). Supplementary Figure 3 illustrates the subsampled age distributions and the sample sizes for normal and tumor groups. After subsampling, we recomputed the average AA in tumor and normal samples, effectively reproducing Figures 2 and Supplementary Figure 1. As shown in Supplemetary Figures 4 and 5, the results remained consistent. We therefore conclude that age does not affect the age acceleration difference between normal and tumor samples in the set of cancers and DNAm clocks.

### 3.3 Cross-Tissue Patterns of Tumor-Associated Clock CpGs

Visualization of clock CpG methylation revealed marked differences between normal and tumor tissues. For example, as illustrated for the Horvath clock in lung cancer (Figure 4A), where the two groups showed distinct methylation patterns across many clock CpGs. We generated similar heatmaps for the other tissues and epigenetic clocks and consistently observed a higher proportion of CpGs with intermediate beta values in tumor samples. In many cases, these CpGs had beta values close to 0 or 1 in normal samples but shifted toward intermediate methylation levels in tumors, which may reflect increased epigenetic heterogeneity or allele-specific methylation, with one allele methylated and the other unmethylated. We next examined whether individual CpGs that discriminated tumor from normal samples were shared across cancer types. The UpSet analysis demonstrated substantial cross-tissue sharing, although the size and composition of the intersections differed among clocks (Figure 4B; Supplementary Figure 6). For EN and BLUP, the largest exact intersections contained CpGs shared across nearly all tissues but excluded thyroid, whereas Skin Horvath and Hannum showed prominent intersections shared across all eight tissues.

**FIGURE 4.**
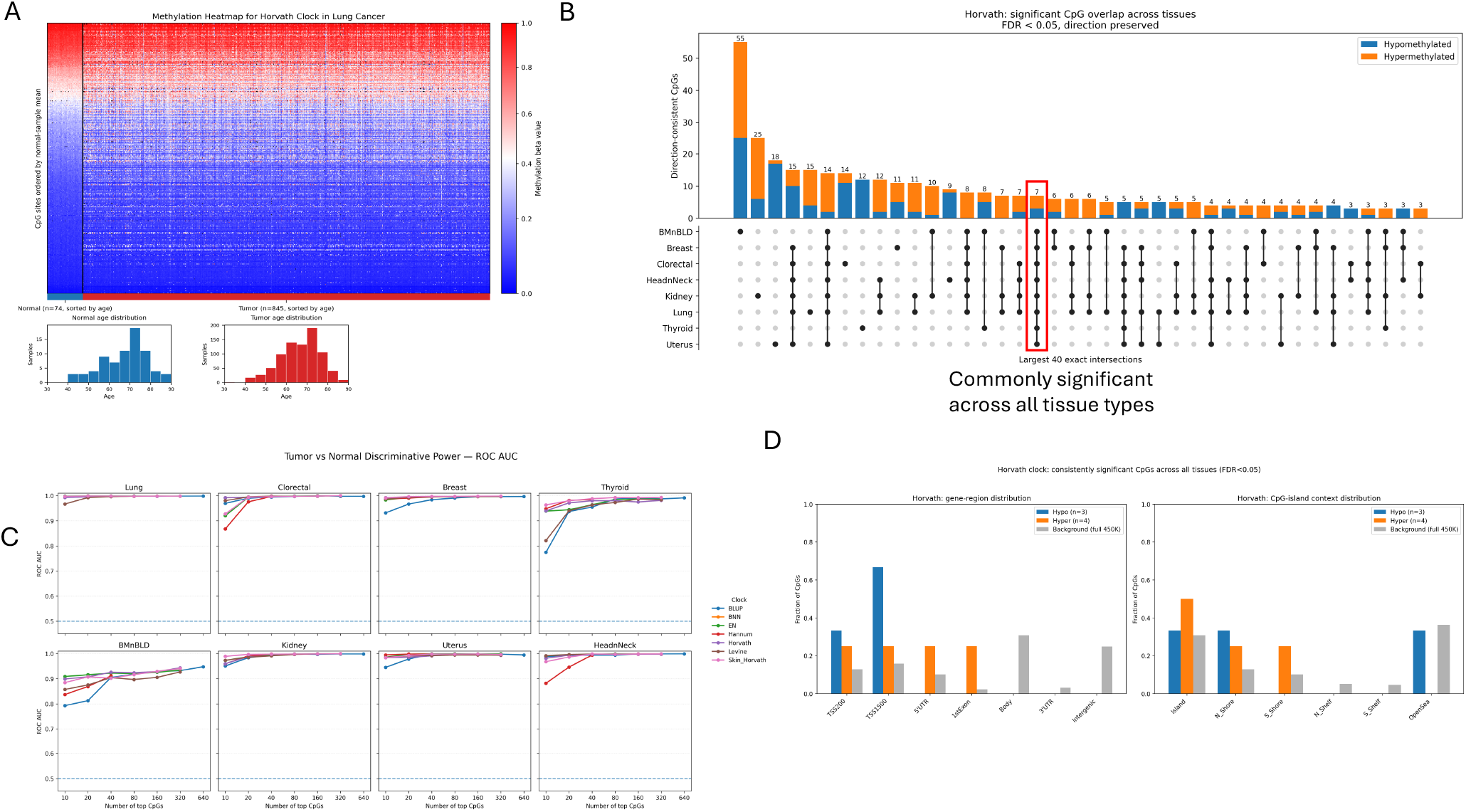
(A) Representative methylation heatmap for CpGs included in the Horvath clock in lung tissue. Normal and tumor samples are shown separately and ordered by increasing age within each group, while CpGs are ordered according to their mean methylation beta value in normal samples. Age distributions for the normal and tumor cohorts are shown below the heatmap. (B) UpSet plot showing the overlap of Horvath clock CpGs significantly associated with tumor versus normal status across the eight tissue types. Bars indicate the number of CpGs in each exact tissue intersection; the highlighted intersection represents CpGs that were significant across all tissues. (C) Genomic annotation of Horvath clock CpGs that were consistently significant across all tissues at FDR < 0.05. Hypomethylated and hypermethylated CpGs are shown according to their gene-region annotation and CpG-island context, with the distribution of all probes on the Illumina HumanMethylation450 array shown as background for comparison. (D) Comparison of the tumor–normal discriminatory power of CpGs from different epigenetic clocks. CpGs were ranked according to their univariate tumor–normal association, and elastic-net logistic regression models were evaluated using increasing numbers of top-ranked CpGs. Curves show cross-validated ROC AUC for each clock and tissue, with the dashed line indicating the random-classification baseline (ROC AUC = 0.5).

Absolute intersection sizes were particularly large for BLUP, reflecting in part the much larger number of CpGs represented by this clock and therefore preventing direct comparison of intersection counts between clocks. When the analysis was restricted to CpGs that were consistently significant across all eight tissues at FDR < 0.05 while preserving the direction of methylation change, tumor-associated hypermethylation predominated. Skin Horvath contained 21 such CpGs (18 hypermethylated and 3 hypomethylated), EN contained 12 (8 hypermethylated and 4 hypomethylated), Horvath, BNN, and Hannum each contained 7 (4 hypermethylated and 3 hypomethylated), and Levine contained 7 (5 hypermethylated and 2 hypomethylated). Genomic annotation of these CpGs showed that consistently hypermethylated sites frequently occurred within or near CpG islands, whereas hypomethylated sites were more broadly distributed across CpG-island contexts. Their distribution across gene-associated regions was more clock-specific, with representation in promoter-proximal regions, gene bodies, untranslated regions, and intergenic regions varying among clocks (Figure 4D; Supplementary Figure 7). Given the relatively small number of consistently significant CpGs for most clocks, these genomic distributions were interpreted descriptively rather than as formal enrichment tests. The supplementary material contains the full cross-clock UpSet and compartment analyses.

### 3.4 Discriminatory Power of Clock-Specific CpG Sets

We next compared how efficiently CpGs from each epigenetic clock captured the methylation differences between tumor and normal tissues. Across most cancer types, strong discrimination was achieved using relatively small numbers of the highest-ranked CpGs (Figure 4C). ROC AUC values were generally above 0.9 with as few as 10 CpGs and approached 1.0 as additional CpGs were included for lung, colorectal, breast, kidney, uterus, and head-and-neck cancers. Differences among clocks were most apparent when only a small number of CpGs were used; for example, BLUP and Levine showed lower initial performance in thyroid, while Hannum showed lower performance with small CpG sets in colorectal and head-and-neck cancers, before converging toward the other clocks as additional CpGs were included. Bone marrow and blood showed the lowest overall discriminatory performance and the clearest separation among clocks, with performance increasing more gradually as additional CpGs were added. The PR AUC analysis showed a similar pattern, with values approaching 1.0 for most solid tumor cohorts and lower, but still substantially above the prevalence-based baseline, performance in bone marrow and blood (Supplementary Figure 8). Overall, performance generally plateaued after approximately 20–80 top-ranked CpGs for most tissues, indicating that a relatively small subset of the CpGs represented in each clock captured much of the methylation signal distinguishing tumor from normal tissue.

### 3.5 Cancer-Related Genes Among Discriminatory CpGs

Gene annotation of CpGs that were consistently discriminatory across all tissues identified several loci with independent evidence of cancer-associated epigenetic dysregulation (Supplimentary table 1). Notably, the hypomethylated DDR1-associated CpGs in the Horvath and BNN clocks are concordant with reported loss of DDR1 methylation in lung cancer [36]. Among the hypermethylated loci, TBX5 has been described as an epigenetically silenced tumor suppressor in colorectal cancer [37], RXFP3 is frequently promoter-hypermethylated in endometrial carcinoma [38], and NDRG2 undergoes methylation-associated silencing in colorectal cancer [39]. Particularly striking was the Skin Horvath TBR1 signal, which mapped to the 3^′^UTR, matching a renal-cancer-associated hypermethylated region reported in the same genomic compartment [40]. Skin Horvath also contained multiple clustered protocadherin genes, consistent with coordinated hypermethylation and long-range epigenetic silencing of this locus in solid tumors [41]. Together, these examples support overlap between age-associated clock CpGs and recurrent cancer-associated epigenetic remodeling.

## 4 Discussion

This study analyzed age acceleration (AA) across eight cancer types using seven epigenetic clocks and 5,528 samples, comparing tumor and normal tissues. Overall, tumor samples showed higher AA than normal tissues, consistent with prior findings that link cancer to accelerated aging. Notably, the uterus cohort was an exception, where normal tissues displayed higher AA. Individual-level analysis confirmed that predicted ages for tumor samples were generally higher than for normal tissues, highlighting the impact of cancer on epigenetic aging. Subsampling confirmed that these results were not affected by age distribution differences. Our study supports AA as a potential biomarker for cancer prognosis and highlights unique cohort-specific patterns that warrant further investigation.

The discriminatory power of clock-specific CpG set analysis was intended to provide a relative comparison of the tumor–normal discriminatory information captured by the CpGs included in different epigenetic clocks, rather than to develop or evaluate a clinically applicable cancer classifier. CpGs were ranked using their tumor-versus-normal associations calculated from the full dataset before cross-validation, which introduces information leakage between feature selection and model evaluation and may lead to optimistic estimates of ROC AUC and PR AUC. Therefore, the reported predictive performance should not be interpreted as an estimate of how these CpG sets would perform for cancer diagnosis in independent patients. A rigorous diagnostic model would require feature selection to be performed exclusively within each training fold, appropriate hyperparameter tuning using nested cross-validation, and validation in an independent external cohort. Here, the classification analysis was used only as a comparative framework to assess how rapidly CpG sets from different clocks capture methylation patterns that distinguish tumor from normal tissue.

