## Supplemental_figures for "Epigenetic Clocks Reveal Age Acceleration and Shared Methylation Remodeling Across Cancers"

#### Supplementary Material

Saleh Sereshki and Stefano Lonardi

### Supplementary Figures

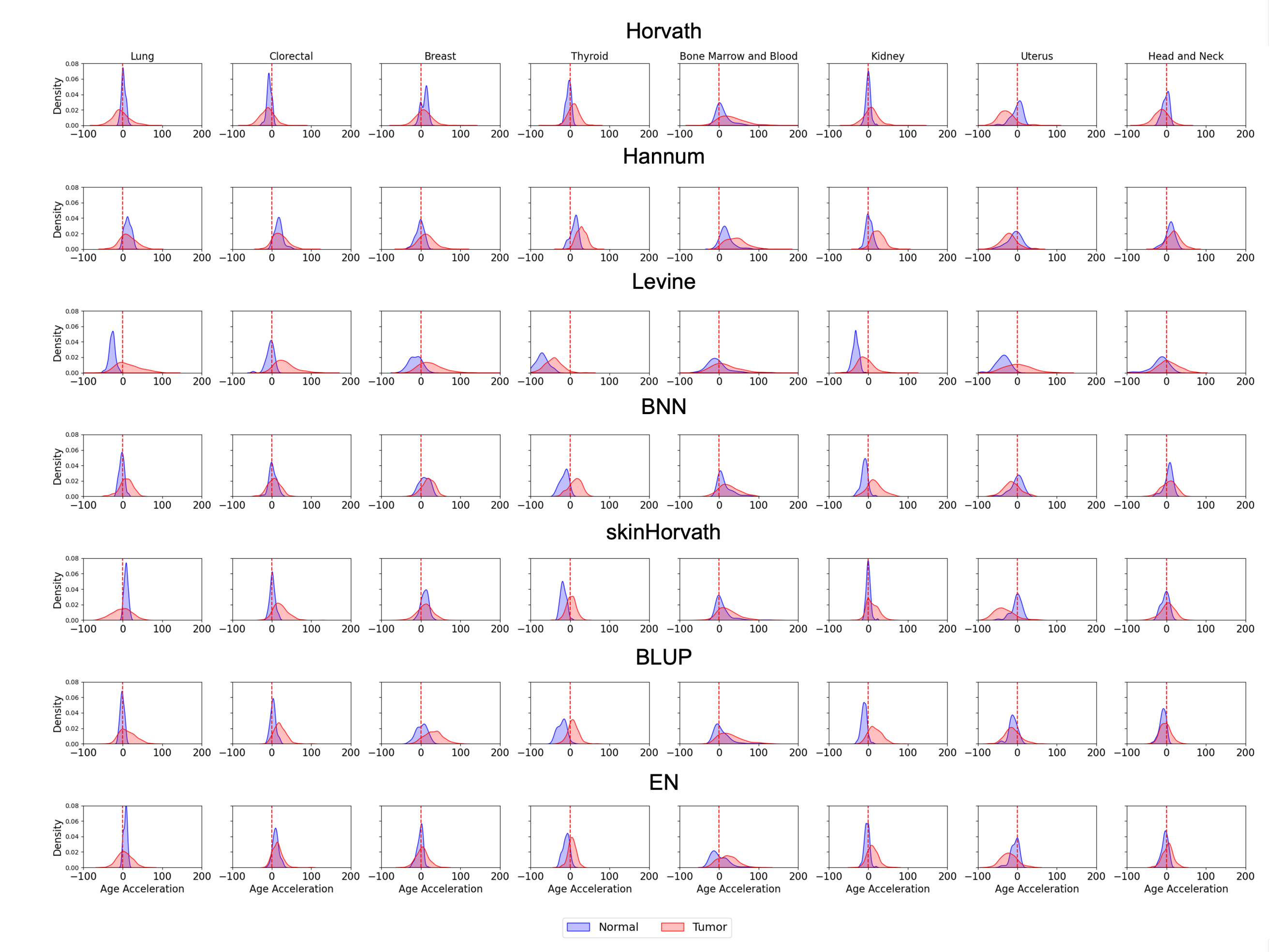


Supplementary Figure 1: Density distributions of age acceleration are shown for normal and tumor samples across eight cancer types and seven epigenetic clocks (Horvath, Hannum, Levine, BNN, Skin Horvath, BLUP, and EN). Rows correspond to epigenetic clocks and columns to cancer types. Normal and tumor samples are shown separately, allowing comparison of shifts in age-acceleration distributions between the two tissue groups. The vertical dashed line indicates zero age acceleration.

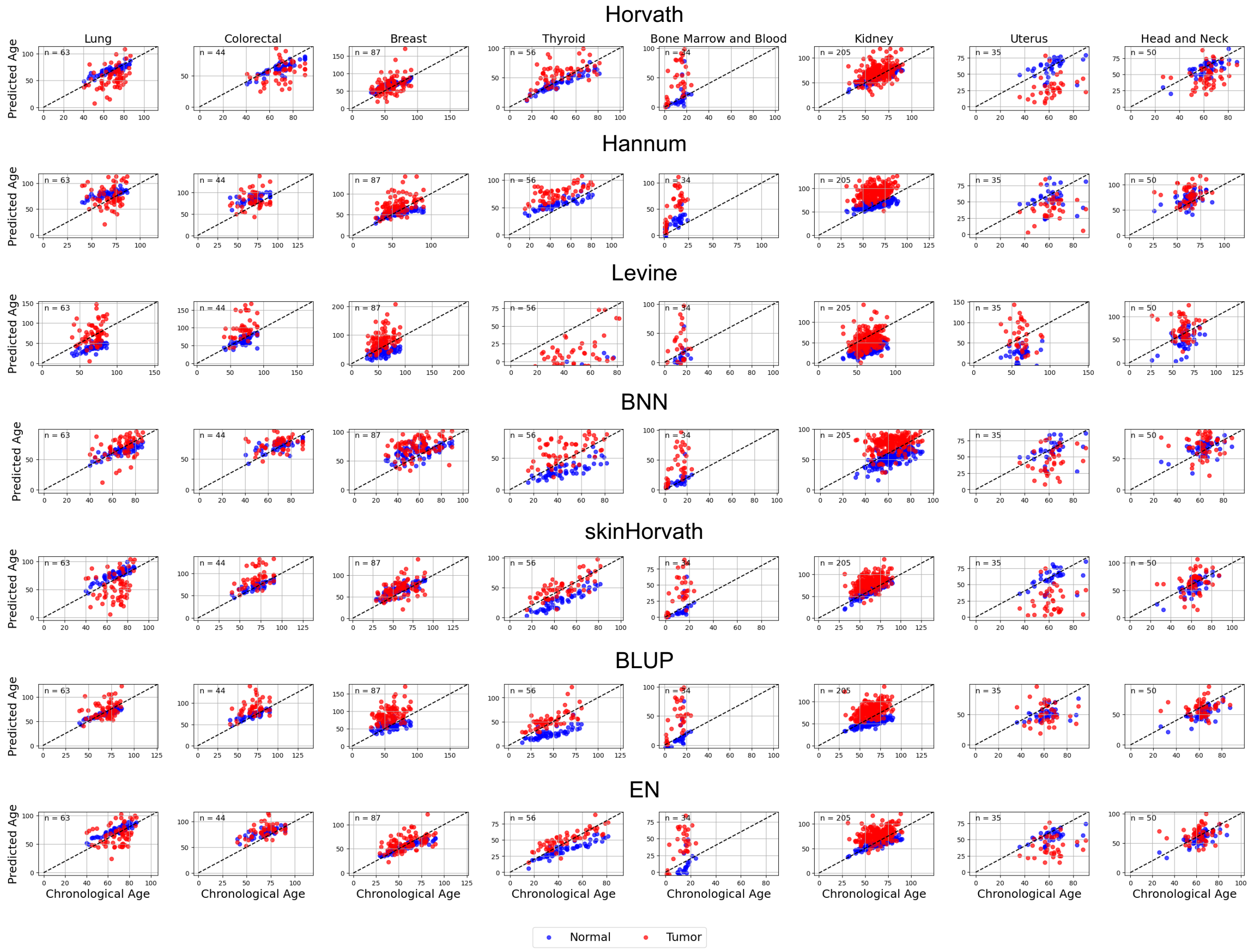


Supplementary Figure 2: Scatter plots compare biological age predicted by seven epigenetic clocks with chronological age across eight cancer types for individuals with one tumor and one normal sample.

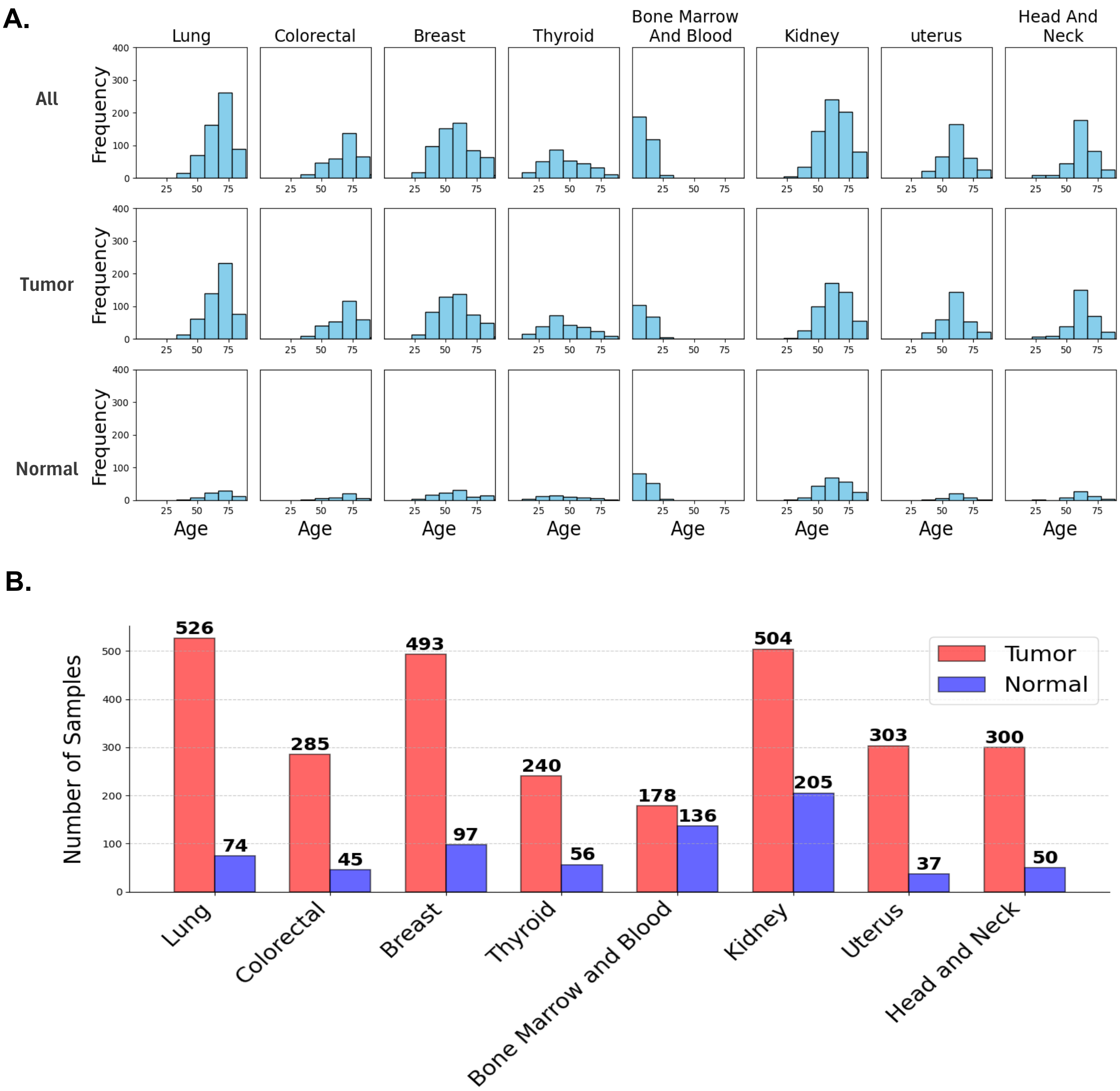


Supplementary Figure 3: Reproduction of the cohort characterization shown in Figure 1 of the main manuscript after age-stratified subsampling

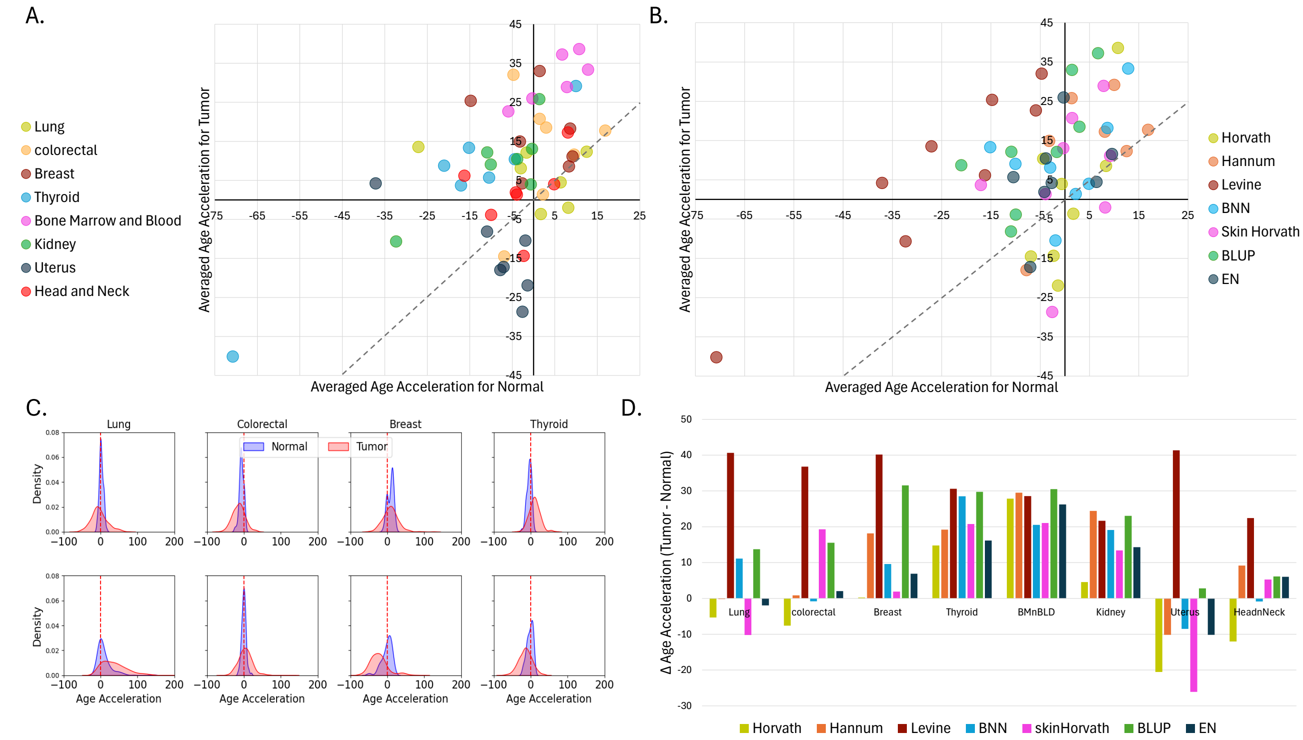


Supplementary Figure 4: Reproduction of the age-acceleration analyses presented in Figure 2 of the main manuscript using the age-stratified subsampled cohorts.

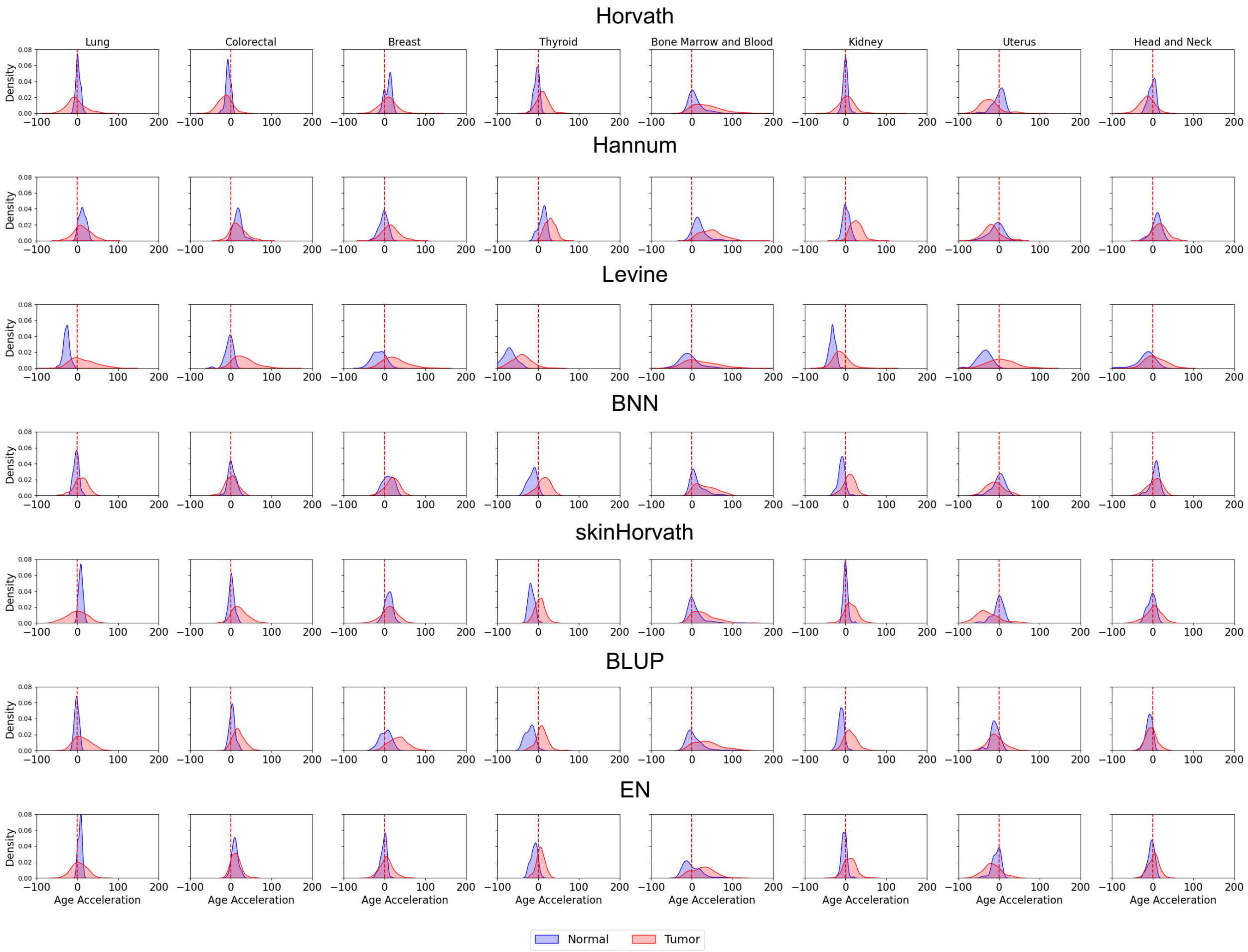


Supplementary Figure 5: Reproduction of Supplementary Figure 1 using the age-stratified subsampled cohorts.

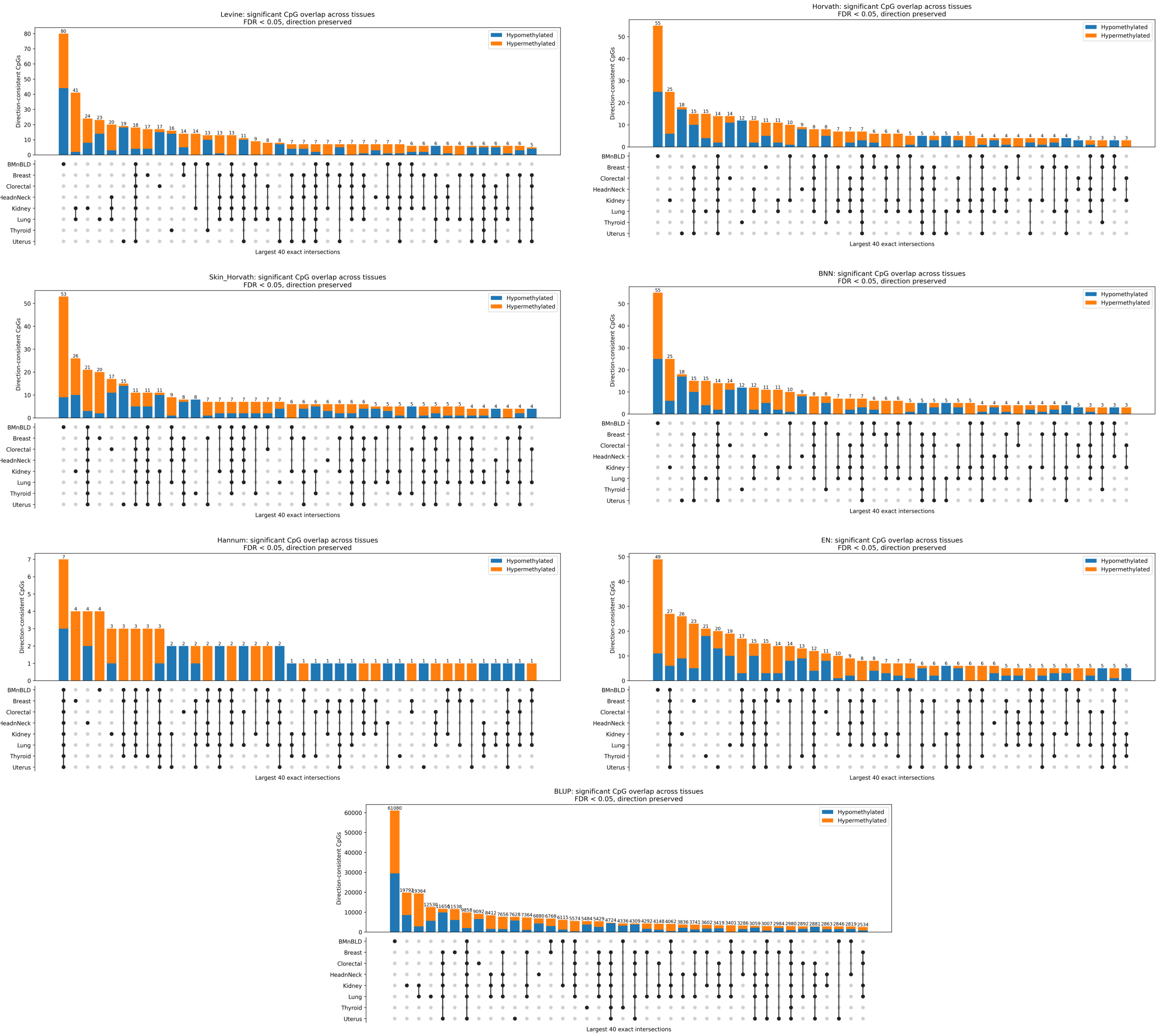


Supplementary Figure 6: UpSet plots summarize the overlap of clock CpGs significantly associated with tumor versus normal status across the eight tissue types at FDR ¡ 0.05 with the direction of methylation change preserved. Bars indicate the number of CpGs belonging to each exact tissue intersection and distinguish CpGs showing tumor-associated hypomethylation from those showing hypermethylation. The matrix below each bar identifies the combination of tissues contributing to that intersection.

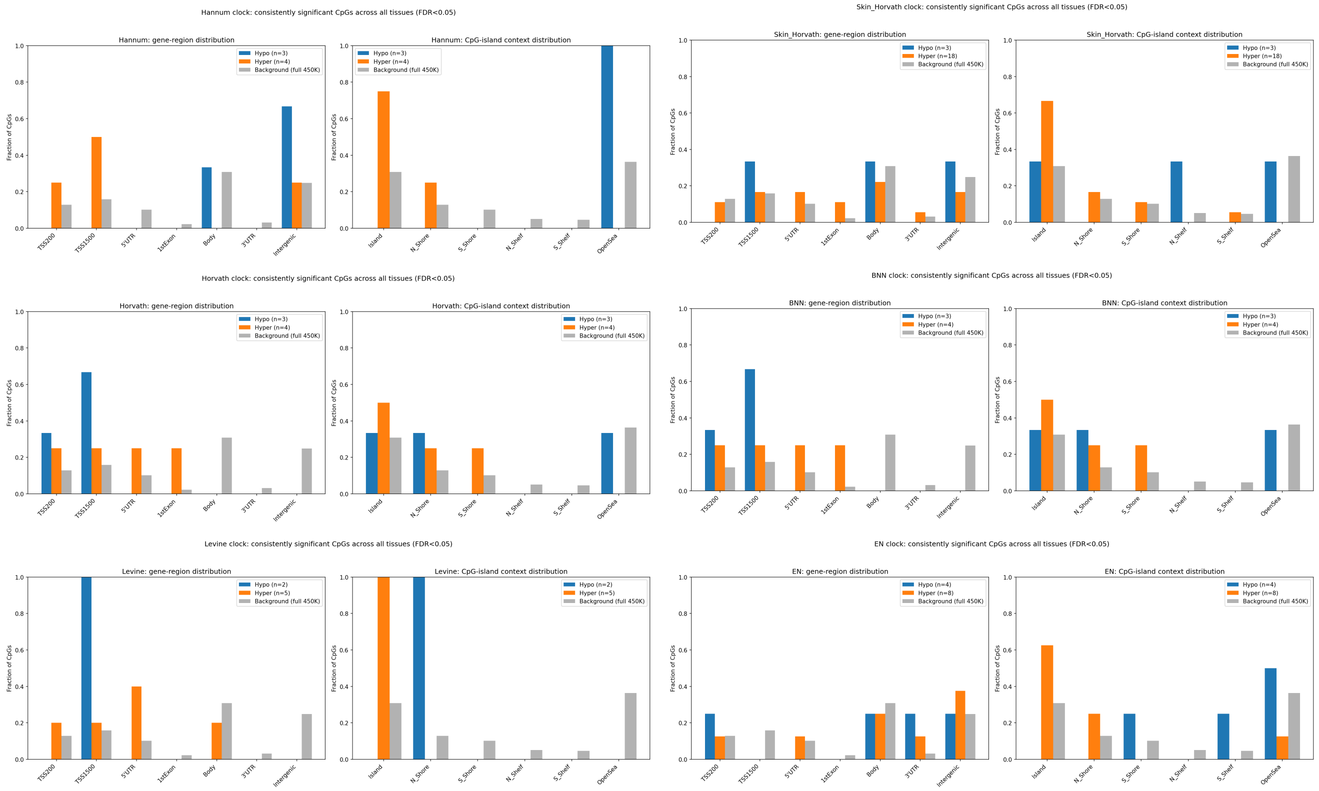


Supplementary Figure 7: Genomic annotation of CpGs that were significantly associated with tumor versus normal status across all eight tissues at FDR $< 0.05$, shown separately for each epigenetic clock. For each clock, the left panel shows the distribution of consistently hypomethylated and hypermethylated CpGs across gene-associated regions, including TSS200, TSS1500, 5UTR, first exon, gene body, 3UTR, and intergenic regions. The right panel shows their distribution according to CpG-island context, including islands, shores, shelves, and open-sea regions. The corresponding distribution of all probes on the Illumina HumanMethylation450 array is shown as a background reference.

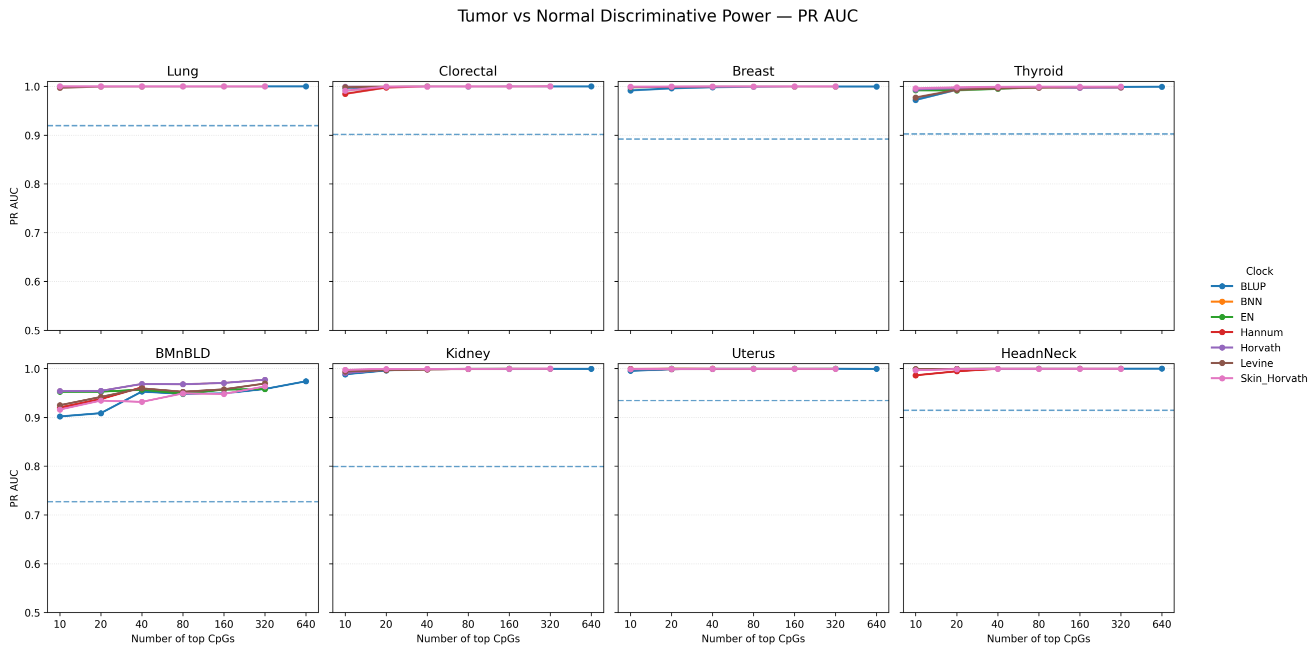


Supplementary Figure 8: PR AUC values are shown for elastic-net logistic regression models constructed using increasing numbers of the most strongly tumor–normal-associated CpGs from each epigenetic clock. Models were evaluated separately across the eight cancer types using the top 10, 20, 40, 80, 160, 320, and 640 CpGs, where available. Each line represents an epigenetic clock. The dashed horizontal line indicates the prevalence-based random-classification PR AUC for the corresponding cancer dataset.
